# Extended-spectrum beta-lactamase (ESBL)-producing *E. coli* in livestock and free-roaming wildlife: a combined phenotyping – whole-genome sequencing One Health approach

**DOI:** 10.1101/2025.04.08.647752

**Authors:** Sheena Conforti, Adamandia Kapopoulou, Claudia Bank, Belinda Köchle, Bahtiyar Yilmaz, Jens Becker

**Affiliations:** Eawag, Swiss Federal Institute of Aquatic Science and Technology, 8600 Dübendorf, Switzerland; Institute of Ecology and Evolution, University of Bern, 3012 Bern, Switzerland; Swiss Institute of Bioinformatics, 1015 Lausanne, Switzerland; Clinic for Ruminants, Vetsuisse-Faculty, University of Bern, 3012 Bern, Switzerland; Department of Visceral Surgery and Medicine, Bern University Hospital, University of Bern, 3010, Bern, Switzerland; Maurice Müller Laboratories, Department for Biomedical Research, University of Bern, 3008, Bern, Switzerland

**Keywords:** environment, selection pressure, antimicrobials, resistance

## Abstract

Antimicrobial resistance (AMR) in Enterobacterales, particularly extended-spectrum beta-lactamase-producing *Escherichia coli* (ESBL-Ec), poses a significant public health concern. Widespread antimicrobial use exerts selection pressure, driving the persistence and spread of resistant bacteria in diverse environments. While ESBL-Ec is well-documented in clinical and agricultural settings, its presence in reservoirs remains poorly understood. This study assessed the prevalence, AMR profiles, and genetic diversity of ESBL-Ec in Swiss dairy cattle and wildlife. Between 2021 and 2023, 775 samples were analysed, including rectal swab samples from dairy cows (n=475), and fecal or cloacal swab samples and wildlife (n=300). ESBL-Ec was detected in 0.6% of dairy cattle and 3.7% of wildlife. Whole-genome sequencing of 54 ESBL-Ec isolates identified 28 sequence types with no overlap between livestock and wildlife, suggesting distinct evolutionary trajectories. Phenotypic testing revealed resistance beyond beta-lactams, notably against aminoglycosides, tetracyclines, and sulfonamides, whereas all isolates remained susceptible to tigecycline and meropenem. Multidrug resistance was prevalent (89%), with bla_CTX-TEM1_ (in dairy cattle), bla_CTX-M55_ and bla_TEM-176_ (in red foxes) widely distributed across Switzerland. Clonal plasmids (IncFIB, IncX1) were detected across hosts and some individuals harbored multiple ESBL-Ec strains. Despite their low prevalence, spatial clustering indicated local persistence in both livestock and wildlife, highlighting potential transmission. The observed presence of clinically relevant ESBL-Ec in wildlife highlights the need for a One Health approach to tackle the dangers of AMR. Our findings contribute to AMR surveillance by providing baseline data on ESBL-Ec reservoirs and informing the design of strategies to mitigate its environmental spread.

## Introduction

Clinically relevant Enterobacterales pose a significant threat to public health and their dissemination has garnered considerable attention in recent decades [1,2]. Of particular concern are isolates carrying extended-spectrum beta-lactamase (ESBL) genes, which confer resistance to a wide range of critically important antimicrobial agents, including extended-spectrum penicillins as well as third- and fourth-generation cephalosporins. These ESBL-encoding genes are often located on mobile genetic elements, facilitating their rapid spread across diverse environments, irrespective of the presence of direct antimicrobial selection pressures.

Over the past decade, in Europe, ESBL-producing *Escherichia coli* (ESBL-Ec) have been detected beyond healthcare settings, notably in livestock, wastewater, and wildlife, underscoring the necessity of a One Health approach to mitigate this public health challenge [3–7]. However, the transmission dynamics of ESBL-Ec remain poorly understood, hindering the ability to predict their spread and persistence. Consequently, regular surveillance, including prevalence assessments, and phenotypic and genotypic characterizations of ESBL-Ec, are important to identify sources of dissemination, track emerging resistance patterns, and inform targeted interventions to mitigate their impact. Discrepancies between genotypic and phenotypic resistance occur, highlighting the complexity of the mechanisms of antimicrobial resistance (AMR) and the importance of investigating resistance at both levels. Previous studies have shown that combining phenotypic and genotypic characterization improves the prediction of AMR resistance patterns and informs more accurate strategies for surveillance and control [8].

Numerous studies on ESBL-Ec in the European Union have focused on specific species or geographic areas [9,10]. In Switzerland, efforts have focused on swine, calves, and poultry, as highlighted in the Swiss Antibiotic Resistance Report, but data on indicator *E. coli* in dairy cattle and wildlife are lacking [11]. Complementing these ongoing surveillance programs is important, as dairy cattle and wildlife differ significantly in terms of antimicrobial selection pressure, habitats, ecological niches, and interactions with humans, representing two distinct potential reservoirs of ESBL-Ec.

This study addresses critical gaps in antimicrobial resistance surveillance by (a) estimating the prevalence of ESBL-producing *E. coli* in dairy cattle and wildlife, (b) characterizing phenotypic co-resistance across antimicrobial classes, and (c) integrating whole-genome sequencing for genotypic resistance characterization. This study reports prevalence, AMR epidemiology, resistance profiles, and clonal dynamics of ESBL-Ec in Swiss dairy cattle and wildlife, offering new data to refine surveillance strategies and inform targeted mitigation efforts.

## Material and Methods

### 2.1. Sample collection

Samples were retrieved from two field studies investigating AMR in Swiss dairy cows and wildlife between 2021 and 2023 [12,13]. Target populations included nationwide dairy cattle, and free-roaming wild mammals and birds from Switzerland and the Principality of Liechtenstein. Both live and recently hunted or deceased animals were sampled. Rectal or cloacal mucosa swabs were collected, and for small birds, fecal samples were obtained after defecation on sterile surfaces. Swabbing was performed using sterile cotton swabs (Copan Transystem, Brescia, Italy; Meus S.r.l., Padova, Italy) inserted into the rectum or cloaca and placed immediately into transport medium suitable for Enterobacteriaceae. Samples were transported to accredited laboratories, either via courier on the day of collection or by overnight mail. Livestock samples were collected by the study team, whereas wildlife samples were mostly obtained through collaboration with wildlife guardians, hunters and ornithologists. Ethical approvals for all sampling procedures were obtained (permits BE76/2021, P38, and BE61/2020/32505).

### 2.2. Isolation and identification of ESBL-Ec

Rectal swabs were streaked onto CHROMagar ESBL chromogenic media (CHROMagar ESBL, Paris, France) without prior dilution the day after swab sample collection, and incubated at 37°C for 24 h. Up to three dark pink to reddish colonies per plate, indicative of *E. coli*, were selected and confirmed by matrix-assisted laser desorption/ionization time-of-flight mass spectrometry (MALDI-TOF M, Bruker, Bremen, Germany). Animals were considered ESBL-Ec-positive if at least one colony from the CHROMagar ESBL plate was confirmed as *E. coli* by MALDI-TOF MS. The number of ESBL-Ec-positive animals was subsequently recorded. Additionally, 16 ESBL-Ec isolates from a prior study, which had been isolated before the start of the present study, were included for whole-genome sequencing and further phenotypic susceptibility testing [14].

### 2.3. Phenotypic susceptibility testing for antimicrobials

Antimicrobial susceptibility testing followed the Clinical and Laboratory Standards Institute (CLSI) disk diffusion guidelines, using 16 drugs from 10 antimicrobial classes. Antimicrobial classes tested were (μg/disc): tetracyclines [tetracycline (30)], sulfonamides [trimethoprim-sulfamethoxazole (1.25/23.75)], penicillins [ampicillin (10), amoxicillin-clavulanic acid (20/10)], cephalosporins [cephazolin (30), cefotaxime (30), cefepime (30), ceftazidime (30)], phenicols [chloramphenicol (30)], aminoglycosides [gentamycin (10), streptomycin (10)], fluoroquinolones [ciprofloxacin (5), nalidixic-acid (30)], glycyclines [tigecycline (15)], macrolides [azithromycin (15)] and carbapenems [meropenem (10)] (Marnes-la-Coquette, France). *E. coli* ATCC 25922 was used as control strain (Remel Inc., Lenexa, USA). Each isolate was cultivated on brain-heart infusion agar plates (BHI) with 5% sheep blood (Bio Rad, Marnes-la-Coquette, France), incubated at 35±2 °C overnight, and standardized to 0.5 McFarland turbidity. The bacterial suspension was spread onto Mueller-Hinton agar plates (MH) using a sterile cotton swab. Afterwards, disks containing antimicrobial drugs were positioned on the inoculated MH agar plates using a disk dispenser (Bio Rad, Marnes-la-Coquette, France). Plates were incubated aerobically at 35±2°C for 16-18 hours, and inhibition zones were measured using a calibrated ruler. Interpretive criteria by CLSI were used to classify isolates as susceptible (S), intermediate (I), or resistant (R) (11). For Tigecycline, interpretive criteria of the European Committee on Antimicrobial susceptibility Testing (EUCAST) were applied since no CLSI breakpoints were available [16]. Intermediate isolates were considered susceptible. Multidrug resistance (MDR) was defined as resistance to at least one drug from three or more antimicrobial classes [17].

### 2.4. DNA extraction and whole-genome sequencing

DNA was extracted using the DNeasy Blood and Tissue kit (Qiagen, Hilden, Germany) following the manufacturer’s instructions, starting from 1 mL of enriched liquid culture in Luria Broth (AppliChem) and eluting in 100 μL Buffer AE. Whole-genome sequencing (WGS) was performed using the LITE Library Prep method and Illumina sequencing on a NextSeq 1000 P2 flow cell (300 bp paired-end) at Earlham Institute, Norwich, UK.

### 2.5. Bioinformatic analyses and isolates characterization

Illumina reads were trimmed using Trimmomatic v0.35 in paired-end mode, with Illumina Nextera adapters removed (parameters: 2:30:10:1). Quality trimming employed a 4 bp sliding window (quality ≥20), and reads <36 bp were discarded. Read quality pre- and post-trimming was assessed with FastQC v0.11.4. Contamination screening was performed with FastQ Screen v0.15.3 mapping reads to seven reference genomes from NCBI: *Acinetobacter baumannii* (ASM863263v1), *Citrobacter freundii* (ASM381234v1), *Escherichia coli* (MG1655), *Enterobacter roggenkampii* (ASM172980v1), *Klebsiella pneumoniae* (ASM24018v2), *Pseudomonas aeruginosa* (ASM676v1), and *Proteus mirabilis* (ASM6996v1). These species were chosen due to their potential to grow on CHROMagar ESBL plates and co-culture with *E. coli*. Reads were assembled de novo using SPAdes v3.15.9, with error correction disabled and the ‘careful’ option enabled. Genome annotation was conducted with Bakta v1.9.1. Sequence types (STs) were identified via the Achtman MLST scheme using MLST v2.23.0 (https://github.com/tseemann/mlst), and phylogenetic groups were assigned using EzClermont v0.6.3. Antibiotic resistance genes (ARGs) were detected with Abricate v1.0.1, blasting against the CARD database (accessed 4th November 2023). ARGs were classified by antibiotic class based on CARD annotations.

### 2.6. Core genome analyses and phylogenetic reconstruction

The pangenome was generated using Roary v3.13.0 to assess genetic diversity, with core genes (present in ≥99% of isolates) aligned by MAFFT for subsequent analysis. Pairwise SNP differences were calculated using an in-house Python script (https://github.com/banklab/ESBL) and summarized in Table S4. A cutoff of 20 SNPs, based on SNP distribution, was used to define clonal isolates (Table S3). Plasmid content was identified via the PlasmidFinder-2.0 using the Enterobacteriales database, with threshold of ≥95% identity and ≥ 60% coverage, and the raw paired-end fastq reads as input.

Phylogenetic analysis was performed using IQ-TREE 2 on 55 sequences, including 54 ESBL-Ec isolates and one outgroup (*Escherichia albertii*, NCBI: ASM1690475v2). The input alignment (2’212’064 sites) contained 86.74% constant sites, 149’328 parsimony-informative sites, and 58’407 distinct site patterns. The best-fit substitution model (GTR+F+R5) was selected using ModelFinder based on the Bayesian Information Criterion. Maximum likelhood was used to infer tree topology, with branch support assessed via Ultrafast Bootstrap Approximation (UFBoot2) with 1’000 replicates. The tree was visualized and annotated in iTOL.

## Results

### 3.1. Study populations and prevalence of ESBL-Ec

A total of 300 fecal samples from wildlife and 475 rectal samples from dairy cows collected from 131 farms were streaked to isolate ESBL-producing *Escherichia coli* (ESBL-Ec). The sampled wildlife population comprised 23 bird species (n=99, 33.3%) and 15 mammal species (n=200, 66.6%), along with one unidentified animal. Red foxes, European roe deer, wild boars, and European badgers were the most frequently represented species (Table S1). The overall prevalence of ESBL-Ec was 0.6% in dairy cows (3 positive animals) and 3.7% in wildlife (11 positive animals, Table 1). Wildlife animals positive for ESBL-Ec included red foxes (n=4), black-headed gulls (n=2), mallard ducks (n=2), wild boar (n=1), crow (n=1), and roe deer (n=1). In total, nine ESBL-Ec isolates were retrieved from cows and 29 from wildlife. The sample of the carrion crow was imported from Austria prior to shipment to the laboratory, with cantonal authorities confirming that no additional permits were required. Sampling locations for dairy farms and wildlife, along with the locations of ESBL-Ec, including those retrieved from a previous study, are shown in Figure 1.

**Figure 1:**
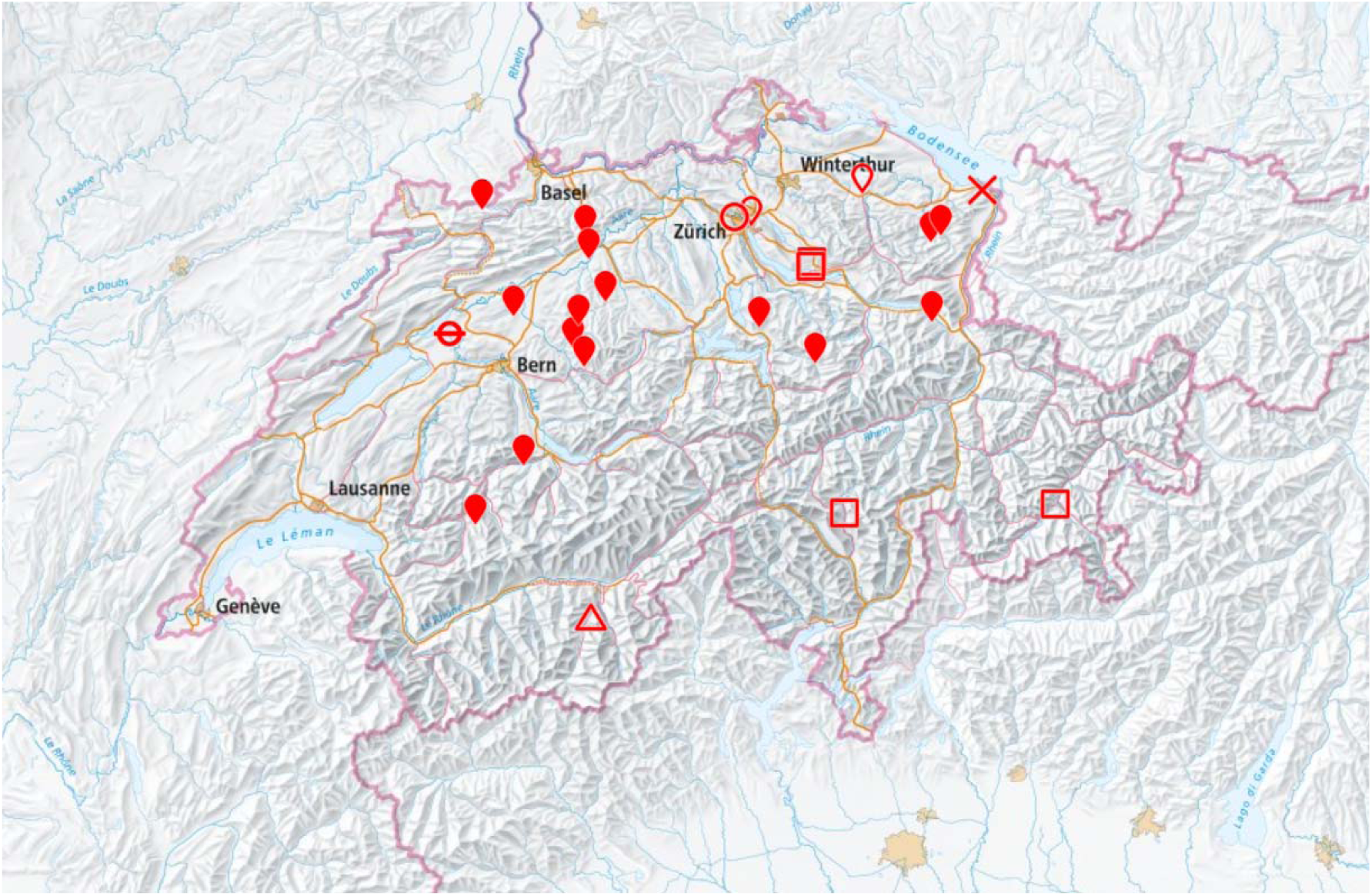
Map of Switzerland with locations of sampled livestock and free-roaming wildlife harboring extended-spectrum beta-lactamase (ESBL)-producing *Escherichia coli*, sampled from 2021-2023 (the map was created using geo.admin.ch, the Swiss federal geoportal) *filled inversed drop: cow, inversed drop: black-headed gull, circle: mallard duck, dashed circle: wild boar, square: red fox, triangle: roe deer, cross: carrion crow*.

**Table 1:**
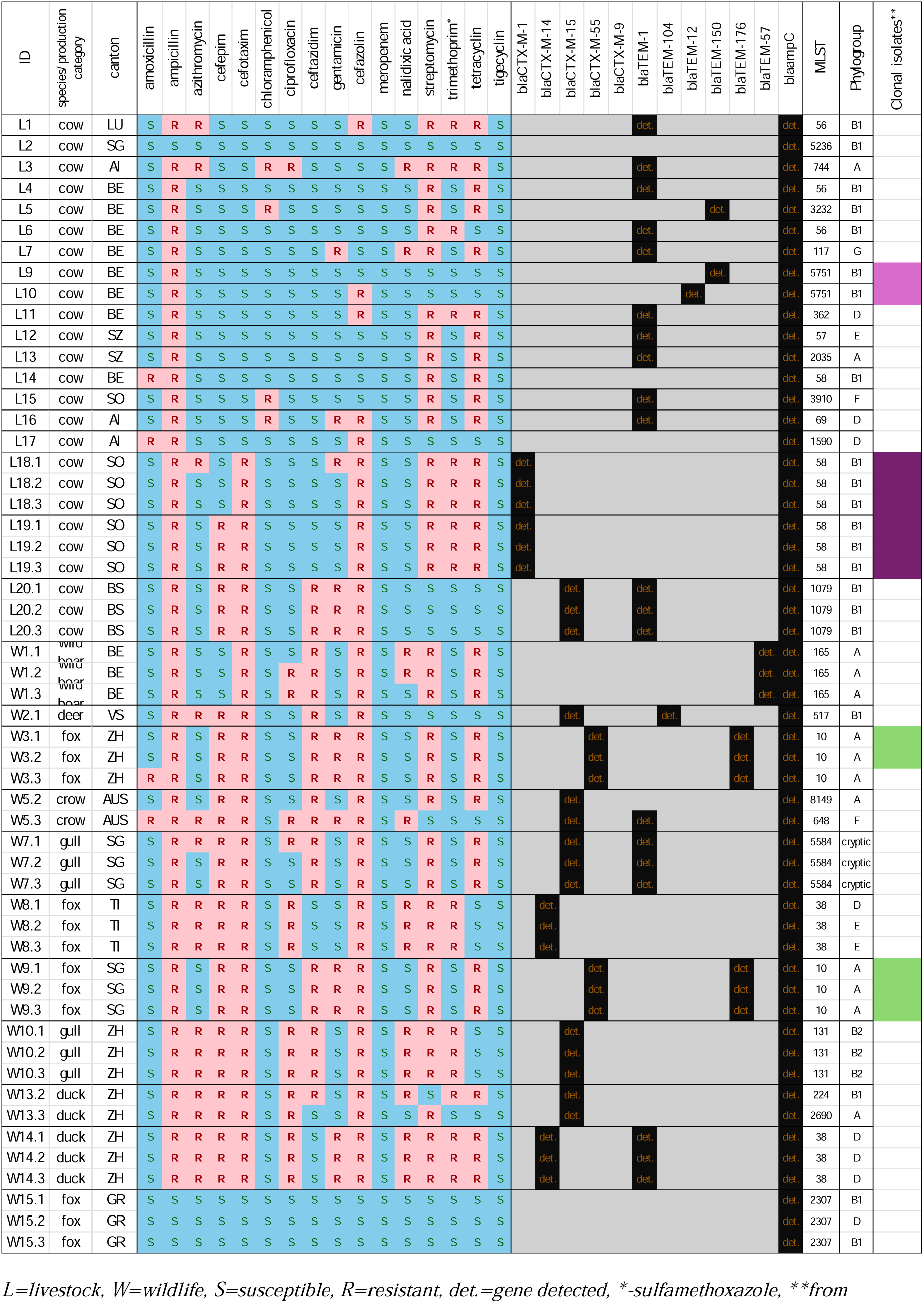
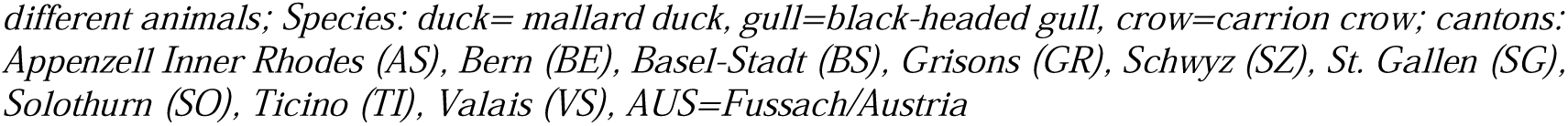
Phenotypic antimicrobial resistance profile, occurrence of genes conferring beta-lactam resistance, and multilocus sequence types (MLST) in 54 *Escherichia coli* isolates from Swiss livestock and wildlife, 2021-2023. Clonal isolates obtained from different individuals are marked with colored bars in the far-right column.

### 3.2. Antimicrobial resistance phenotypes

Phenotyping revealed that 93% (n=50) of the ESBL-Ec isolates exhibited resistance to ampicillin, 7% (n=4) to amoxicillin, 74% (n=40) to cefazolin, 65% (n=35) to the 3^rd^ generation cephalosporin cefotaxime, 41% (n=22) to ceftazidime, and 54% (n=29) to the 4^th^ generation cephalosporin cefepime (Fig. 1). Resistance to other antimicrobial agents included streptomycin (76%, n=41), tetracycline (63%, n=34), trimethoprim-sulfamethoxazole (37%, n=20), ciprofloxacin and nalidixic acid (30%, n=16), gentamycin (30%, n=16) and chloramphenicol (7%, n=4) resistance. No phenotypic resistance was observed to tigecycline and meropenem. Overall, 89% of ESBL-Ec isolates exhibited a multidrug-resistant (MDR) phenotype.

### 3.3 Antimicrobial resistance genotypes

Among the sequenced ESBL-Ec, 89% (n=48) harbored at least one beta-lactamase gene (Table 1). The chromosomally encoded ampC beta-lactamase gene was detected in all isolates. The most frequent gene was bla_TEM-1_, detected in 37% (n=20) of ESBL-Ec, followed by bla_CTX-M-15_, detected in 26% (n=14) of ESBL-Ec. Overall, four different bla_CTX-M_ variants and six different bla_TEM_ variants were detected (Table 1). In decreasing order, the most frequent bla_CTX_ genes were bla_CTX-M-15_ (25.9%, n=14), bla_CTX-M-55_ (11.1%, n=6), bla_CTX-M-14_ (11.1%, n=6), and bla_CTX-M-1_ (11.1%, n=6). Among bla_TEM_ variants, bla_TEM-1_ occurred most frequently (37.0%, n=20), followed by bla_TEM-176_ (11.1%, n=6), bla_TEM_-150 (3.7%, n=2), and bla_TEM-104/-12/-57_ (1.9%, n=1 each). Additionally, 11 genes conferring resistance to aminoglycosides, 5 to fluoroquinolones, 3 to sulfonamides/diaminopyrimidines, 4 to tetracycline, 6 to peptide antimicrobials, 2 to macrolides, and 1 to lincosamides were detected (Figure S2). The total phenotypic and genotypic concordance (i.e., isolates either resistant with ARGs or susceptible without ARGs) was highest for tetracyclines (98%, n=53), followed by beta-lactams (93%, n=50), amphenicols (87%, n=47), aminoglycosides (85%, n=46), sulfonamides/diaminopyrimidines (70%, n=38), and fluoroquinolones (30%, n=16) (Figure 2, Table 2). Notably, it was more frequent for a strain to be phenotypically susceptible but carry a resistance gene than to be phenotypically resistant without a corresponding ARG, except for tetracyclines and amphenicols (Figure 2, Table 2).

**Figure 2:**
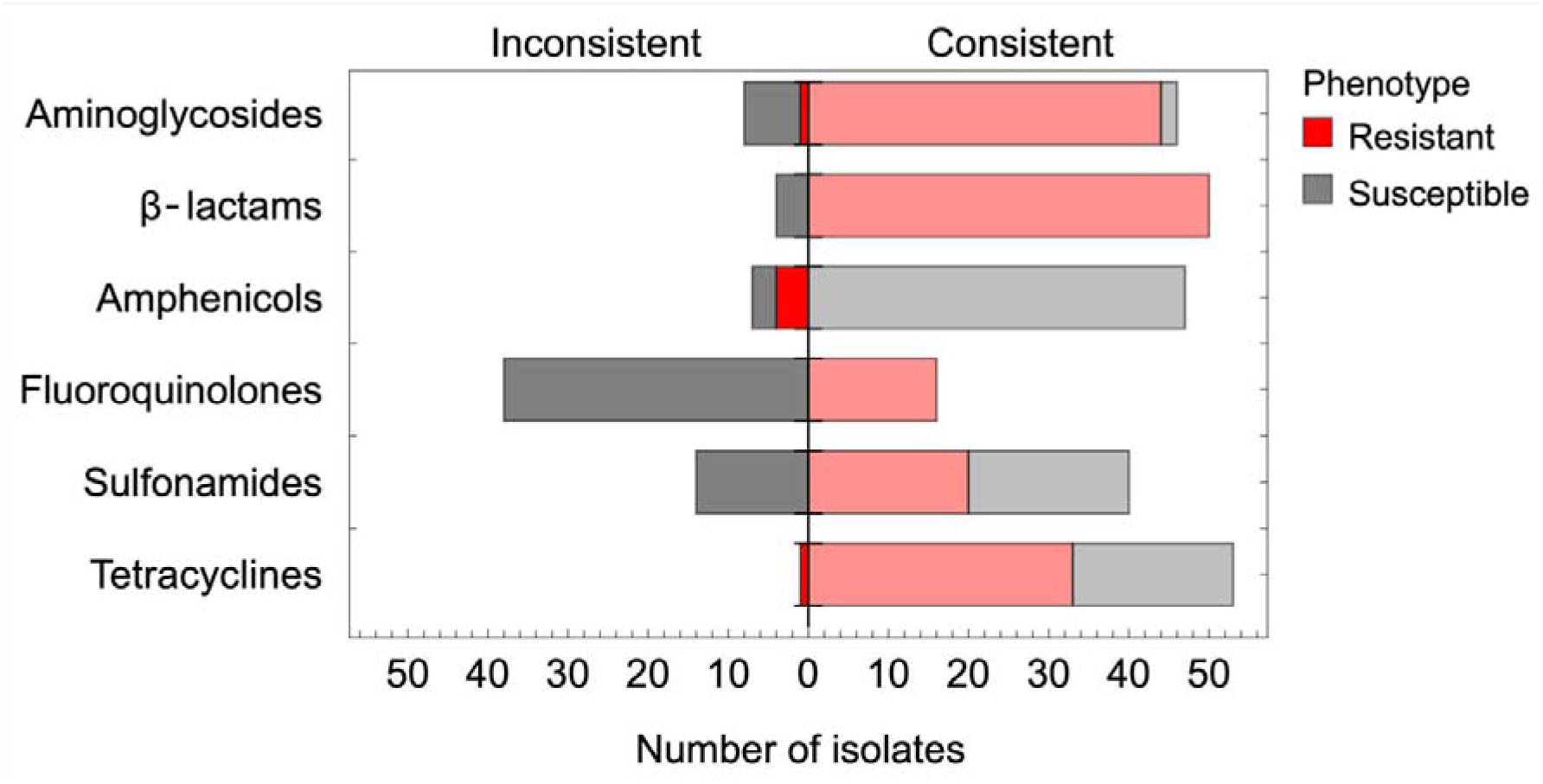
Concordance between phenotypic resistance and antimicrobial resistance gene (ARG) detection across six antibiotic classes. Bars represent the number of isolates with consistent (right of the center line) or inconsistent (left of the center line) results. Consistent results include resistant isolates with the corresponding ARGs (red) and susceptible isolates without the corresponding ARGs (grey). Inconsistent results include resistant isolates without ARGs (red) and susceptible isolates with ARGs (grey).

**Table 2:**
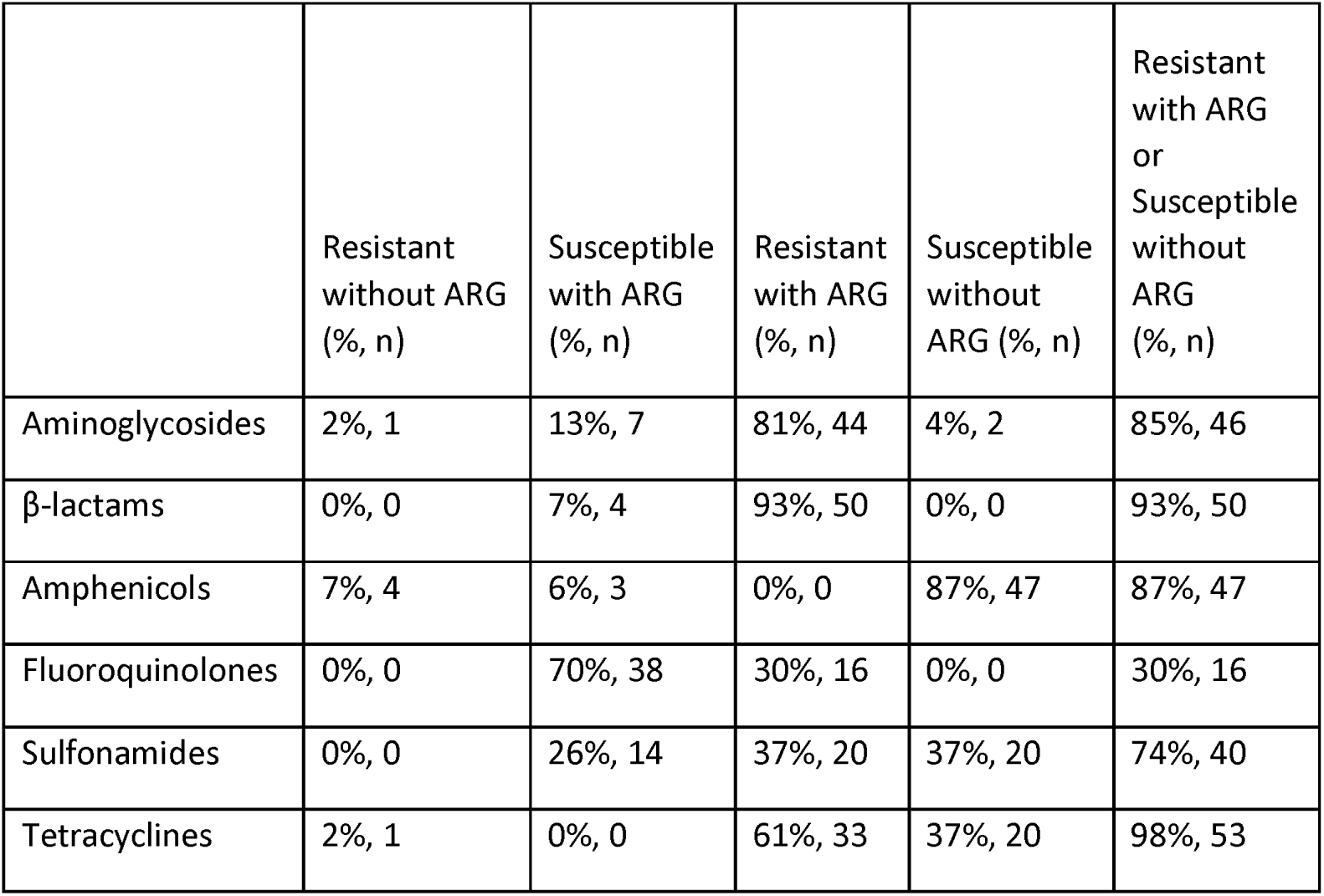
Phenotypic and genotypic concordance for antimicrobial resistance among ESBL-producing *E. coli* isolates, grouped by antimicrobial class. The table presents the percentage and number of isolates classified into five categories: “Resistant without ARG” (phenotypic resistance in the absence of known resistance genes), “Susceptible with ARG” (presence of resistance genes despite phenotypic susceptibility), “Resistant with ARG” (phenotypic resistance with corresponding resistance genes), “Susceptible without ARG” (phenotypic susceptibility with no detected resistance genes), and “Resistant with ARG or Susceptible without ARG” (combined concordance category, representing isolates with matching phenotypic and genotypic resistance or susceptibility). Percentages (%) and absolute numbers (n) are provided for each category

|  | Resistant without ARG<br>(%, n) | Susceptible with ARG<br>(%, n) | Resistant with ARG<br>(%, n) | Susceptible without ARG<br>(%, n) | Resistant with ARG or<br>Susceptible without ARG<br>(%, n) |
| --- | --- | --- | --- | --- | --- |
| Aminoglycosides | 2%, 1 | 13%, 7 | 81%, 44 | 4%, 2 | 85%, 46 |
| β-lactams | 0%, 0 | 7%, 4 | 93%, 50 | 0%, 0 | 93%, 50 |
| Amphenicols | 7%, 4 | 6%, 3 | 0%, 0 | 87%, 47 | 87%, 47 |
| Fluoroquinolones | 0%, 0 | 70%, 38 | 30%, 16 | 0%, 0 | 30%, 16 |
| Sulfonamides | 0%, 0 | 26%, 14 | 37%, 20 | 37%, 20 | 74%, 40 |
| Tetracyclines | 2%, 1 | 0%, 0 | 61%, 33 | 37%, 20 | 98%, 53 |

### 3.4 Phlyogenetic distribution of ESBL-*E. coli* and clonal relationship

The maximum-likelihood phylogenetic tree constructed from a core gene alignment of 54 ESBL-Ec isolates revealed diverse clustering across eight phylogroups, with the most frequent being B1 (39%, n=21), followed by A (24%, n=13), D (15%, n=8), and B2, E and cryptic (6%, n=3 each) (Figure 3). Phylogroup B2 was exclusively observed in gull isolates and corresponded to ST131, a lineage known for its multidrug-resistant phenotype. Among the isolates, 28 unique sequence types (STs) were identified, with the most frequent being ST58 (13%, n=7), followed by ST10 and ST38 (11%, n=6 each), and ST56, ST131, ST165 (6%, n=3 each). Wildlife isolates displayed diversity in STs, with ST38 observed in foxes and ducks and ST10 in foxes, ST131 in gulls, and ST165 in wild boars. ST165 was exclusively detected in wild boars, and ST131 was restricted to gulls. Livestock isolates clustered more narrowly, with ST58 being the predominant sequence type and exclusively found in cattle.

**Figure 3:**
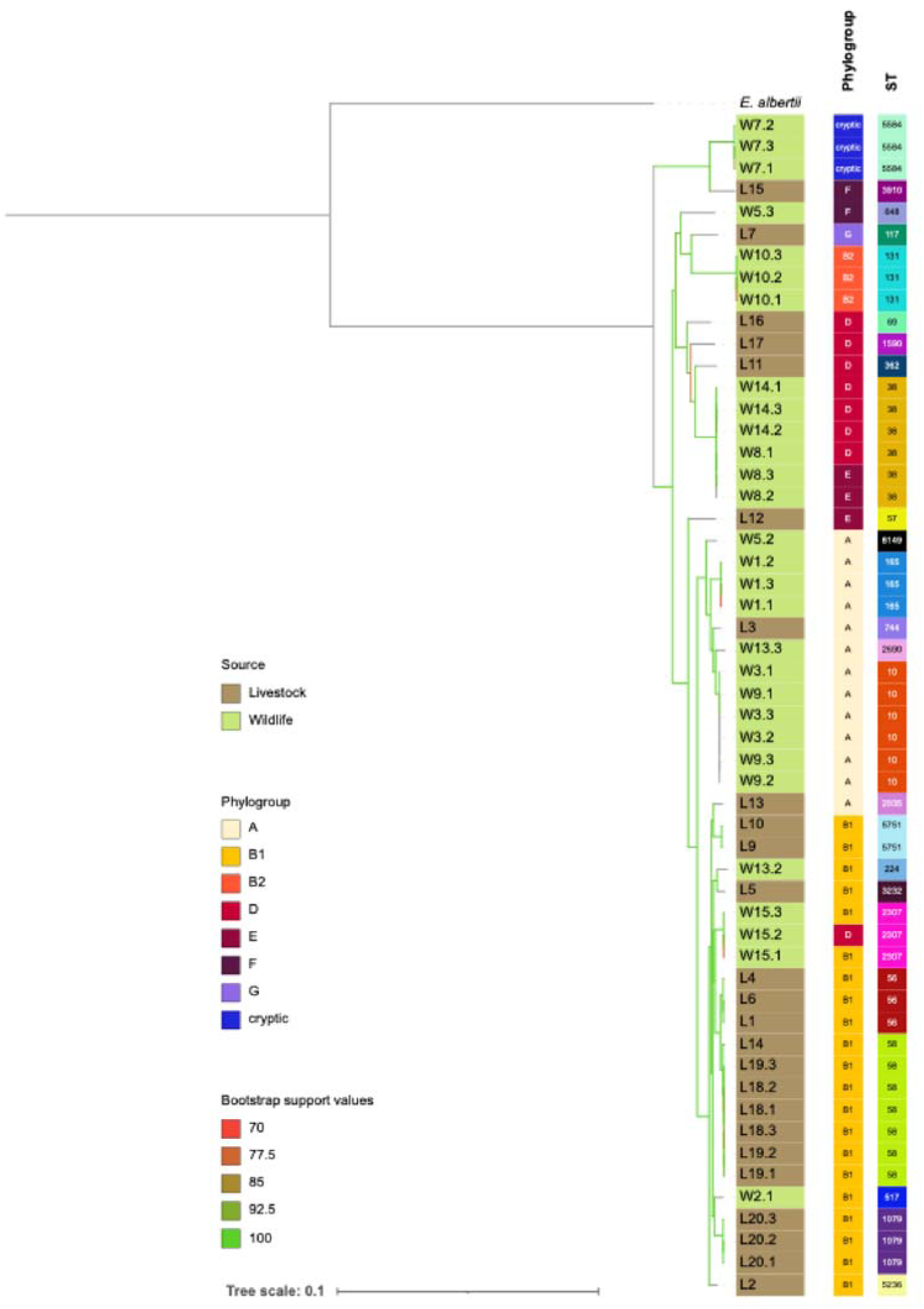
Phylogenetic tree of 54 extended-spectrum beta-lactamase (ESBL)-producing *Escherichia coli* isolates from Swiss livestock and free-roaming wildlife, collected between 2021 and 2023, along with one outgroup (*Escherichia albertii*). The tree illustrates the distribution of phylogroups, sequence types (STs), and sources of collection. Sources are color-coded: brown for livestock and green for wildlife. Phylogroups (A, B1, B2, D, E, F, G, and cryptic) are highlighted with distinct colors. Sequence types (STs) are annotated, with notable STs explicitly labeled. The tree was constructed using IQ-TREE with automatic model selection, and branch lengths represent substitutions per site, as indicated by the scale bar. Bootstrap support values, ranging from 70% (red) to 100% (green), are represented by the color gradient of the branches, with no values below 70% observed.

The phylogenetic tree shows a structured distribution of isolates by source, with some degree of intermixing (Figure 3). Wildlife-derived isolates (green) predominantly cluster in the upper part of the tree, primarily within phylogroups A, D, and cryptic. In contrast, livestock-derived isolates (brown) are more concentrated in the lower clades, particularly within phylogroup B1. Notable exceptions include a wildlife-derived isolate (W2.1, deer) clustering within a primarily livestock-associated clade in phylogroup B1. Similarly, livestock-derived isolates L3 and L13 in phylogroup A cluster within a clade that is predominantly associated with wildlife, reflecting overlapping genetic diversity between the two compartments.

Genetic similarity indicated genetic clusters in both livestock and wildlife resulting in a bimodal distribution of SNP differences between isolates (Table S3). Livestock isolates exhibited either few SNP differences or >400 (mean: 40178 SNPs), and wildlife isolates had either <20 SNP differences or >37 (mean: 46900 SNPs) (Table S4). Clonal relationships were identified among several isolates from different individuals. For example, livestock isolates L9 and L10 had just 2 SNPs differences, whereas isolates L18.1-3 and L19.1-3 differed by 0-8 SNPs (Table S4). Similarly, wildlife isolates W3.1-3 and W9.1-3, all from red foxes in Zurich, demonstrated high degrees of genetic similarity: W3.1 and W9.1 were identical (0 SNPs), and also comparisons between W3.1 and W9.2 (19 SNPs) and W3.1 and W9.3 (16 SNPs) indicated close relatedness (Table S4). In contrast, SNP differences between W3.3 and the W9 isolates were larger (105–123 SNPs), suggesting greater genetic divergence.

## Discussion

This study investigated the prevalence, phenotypic and genotypic antimicrobial resistance profiles, and phylogenetic distribution of extended-spectrum beta-lactamase (ESBL)-producing *Escherichia coli* (ESBL-Ec) in dairy cattle and wildlife in Switzerland from samples collected between 2021 and 2023. The prevalence of ESBL-Ec was 0.6% in dairy cattle and 3.7% in wildlife, with isolates detected across multiple species. Concordance between phenotypic and genotypic resistance was highest for tetracyclines (98%, n=53) and beta-lactams (93%, n=50), but non-concordance occurred where isolates were phenotypically susceptible despite harboring resistance genes or vice versa, especially for fluoroquinolones and sulfonamides. Whole-genome sequencing revealed high genetic diversity between the isolates. Twenty-eight unique sequence types (STs) were identified, with phylogroup B1 being the most frequent (39%, n=21). Close genetic relationships were observed within and between animals, highlighting potential transmission dynamics. The presence of multidrug-resistant lineages such as ST131 in gulls and ST38 in foxes and ducks underscores the public health threat posed by bacteria that reside in environmental reservoirs.

The prevalence of ESBL-Ec in wildlife (3.7%) aligns with studies from Switzerland, Europe, and overseas, where percentages ranged from 1.1% to 10% [6,18–20]. However, higher rates (52%-67%) previously reported from peridomestic wildlife highlight regional and ecological variability of ESBL-Ec prevalence [21,22]. Based on comparison with a Swiss wildlife report from 2011, ESBL-Ec prevalence in wild ungulates has likely remained stable, as our study detected one positive isolate (corresponding to 1.4%) compared to 0.4% reported previously [23]. For Swiss wild birds, a similar prevalence of 1.7% was reported in 2019, aligning with our findings of 1.6% [24]. The detection of ESBL-Ec in wildlife is expected given the frequent use of antimicrobials in livestock, potential direct and/or indirect contact on pastures, and farm effluents which are dispersed onto land without prior treatment, potentially contributing to environmental contamination. Notably, our study design did not allow us to determine whether wildlife serves as a transient host or long-term reservoir for ESBL-Ec.

This study found an ESBL-Ec prevalence of 0.6% in livestock, which is much lower than numbers reported from overseas [21,25–27] and from Switzerland. For example, one Swiss study found a 13.7% prevalence in cattle feces, though this was based on a smaller sample size and used an enrichment step before culturing, which increases detection sensitivity [28]. In contrast, our study applied direct plating without enrichment. Additionally, routine surveillance of veal calves at slaughterhouses in Switzerland has reported ESBL/ampC-Ec prevalence around 30% [29] but these studies used cecal contents collected post-mortem, which may differ microbiologically from rectal swabs. These methodological variations — sample matrix (cecal content vs. feces vs. rectal swabs), enrichment use, animal age group, and sampling context (routine surveillance vs. research study) — complicate direct comparisons and may partly explain the lower prevalence observed here.

We found that multidrug resistance was a prominent feature among the ESBL-Ec isolates, with 89% exhibiting resistance to at least one antimicrobial drug from three or more distinct classes. Besides beta-lactams, phenotypic resistance was most frequent for aminoglycosides, tetracyclines, and sulfonamides, whereas no resistance was detected against tigecycline or meropenem, indicating that these clinically relevant antimicrobials remain effective. The high MDR prevalence in both livestock and wildlife raise concerns about potential transmission of multiple resistance types to other pathogenic bacteria via horizontal gene transfer.

Genetic analysis revealed a diverse range of antimicrobial resistance genes, with bla_TEM-1_ (37%, n=20) and bla_CTX-M-15_ (26%, n=14) as the most prevalent beta-lactamases. Additional resistance determinants for aminoglycosides, fluoroquinolones, sulfonamides/diaminopyrimidines, tetracyclines, peptide antimicrobials, macrolides, and lincosamides were detected. The co-occurrence of multiple resistance genes across unrelated antimicrobial classes may reflect horizontal gene transfer via mobile genetic elements, though independent acquisition cannot be ruled out. [46]. The detection of IncFIB and Inc11-I plasmids in isolates from dairy cattle barns, wild mammals, and birds suggests that these plasmid types may play a role in the dissemination of resistance traits across different environments. Our findings call for in-depth studies of transmission routes of bacteria and plasmids, particularly given the spatiotemporal variability associated with the sampled free-roaming wildlife. Future research should assess whether the spread of AMR via bacteria and/or via plasmids can be mitigated and identify effective intervention strategies.

We found a notable level of non-concordance between phenotypic and genotypic resistance. Isolates were more frequently phenotypically susceptible despite carrying resistance genes than phenotypically resistant without corresponding genes. The non-susceptibility of carriers of resistance genes can potentially be attributed to gene expression variability, differences in promoter activity, or the presence of non-functional resistance genes. Conversely, phenotypic resistance without a detectable gene might indicate alternative and potentially unknown resistance mechanisms, such as via efflux pumps or mutations in target sites. This highlights a limitation of gene-based screening approaches, as they may not fully capture the phenotypic resistance landscape, particularly for resistance mechanisms not associated with well-characterized genes.

Spatial proximity was an important factor related to ESBL-Ec genetic similarity. In wildlife, fox isolates W3.1-3 and W9.1-3, collected 2 km apart at the Zurich-St. Gallen border over a five-week interval, were found to be completely clonal, including their plasmids, suggesting local persistence and potential transmission. In livestock, clonal isolates were frequently detected within the same barns. Dairy cows from tie-stall barns, including isolates L9 and L10 from cows tied at opposite ends of a barn and L18.1-3 and L19.1-3 from another barn under similar conditions, carried genetically similar isolates with less than 10 SNPs difference. The close genetic relatedness of the isolates suggests that confined spaces facilitate clonal spread of ESBL-Ec independent of antibiotic treatment in livestock. Future studies are needed to quantify the duration for which ESBL-Ec persists in barns after cessation of antibiotic treatment.

At a larger scale, phylogenetic analysis revealed some intermixing between livestock and wildlife isolates, with wildlife isolate W2.1 clustering in phylogroup B1, predominantly associated with livestock, and livestock isolates L3 and L13 clustering in phylogroup A, commonly linked to wildlife. However, no sequence type (ST) was shared between livestock and wildlife, suggesting ecological and behavioral barriers limiting direct transmission. The predominance of phylogroup B1 and the exclusive detection of ST58 in cattle indicate livestock as potential reservoirs for specific lineages, whereas ST38 detection in foxes and ducks highlights its adaptability and potential spread across wildlife species.

In Switzerland, *E. coli* ST131 has been repeatedly detected in water and wastewater [7,30] and has spread globally since the early 2000s, causing severe bloodstream and urinary tract infections in humans [31]. Its association with water aquatic environments is well documented [32–34]. Our detection of ST131 in one sample from a gull, a species adapted to such habitats, aligns with reports of ST131 from fish, highlighting its environmental persistence and potential spread across a broad host range. Like ST131, ST38 is linked to MDR and is of global concern [34]. ST38 has been detected in various Swiss environments, including surface water, fish, vegetables, poultry, poultry meat, and healthy humans [35,36]. Our detection of ESBL-Ec ST38 in a duck north of the Alps (phylogroup D) and in a fox from Ticino in southern Switzerland (phylogroups D and E) supports its adaptability to a broad host range and suggests widespread circulation in wildlife across the geographical barrier posed by the Alpine range. This study has several limitations. First, the low ESBL-Ec prevalence in livestock may stem from direct streaking without enrichment, reducing sensitivity for low-abundance bacteria. Second, disk diffusion testing, while standardized, lacks MIC values, limiting insight into resistance levels. Third, whole-genome sequencing, though informative, does not capture all mobile genetic elements or plasmids within one ESBL-Ec isolate. Finally, the cross-sectional design prevents assessing temporal persistence. Longitudinal studies are needed to evaluate clonal stability and the role of livestock and wildlife in sustaining resistance.

In conclusion, this study confirms the presence of ESBL-Ec in both livestock and wildlife across Swiss environments (2021-2023), with evidence of clonal dissemination within each host group. Between host groups, however, ESBL-Ec showed distinct evolutionary trajectories and no shared sequence types, although phylogenetic patterns suggest occasional transmission between wildlife and livestock over longer timescales. The high prevalence of MDR is particularly concerning because it indicates that most frequently, multiple resistance types are maintained and transmitted jointly. Given their occurrence even in remote environments, the results of this study call for the development of targeted strategies and study designs to infer transmission routes and probabilities, the duration over which MDR is maintained, and the selection pressures (or absence thereof) allowing for its maintenance.

## Supporting information

All Supplemental Tables

